# Maturation of prefrontal input to dorsal raphe nucleus increases behavioral persistence in mice

**DOI:** 10.1101/2022.01.01.474690

**Authors:** Nicolas Gutierrez-Castellanos, Dario Sarra, Beatriz S. Godinho, Zachary F. Mainen

## Abstract

The ability to persist towards a desired objective is a fundamental aspect of behavioral control whose impairment is implicated in several behavioral disorders. One of the prominent features of behavioral persistence is that its maturation occurs relatively late in development. This is presumed to echo the developmental time course of a corresponding circuit within late-maturing parts of the brain, such as the prefrontal cortex, but the specific identity of the responsible circuits is unknown. Here, we describe the maturation of the projection from layer 5 neurons of the prefrontal cortex to the dorsal raphe nucleus in mice. We show using pathway-specific optogenetic stimulation that this connection undergoes a dramatic increase in synaptic potency between postnatal weeks 3 and 8, corresponding to the transition from juvenile to adult. We then show that this period corresponds to an increase in the behavioral persistence that mice exhibit in a foraging task. Finally, we use genetic targeting to selectively ablate this pathway in adulthood and show that mice revert to a behavioral phenotype similar to juveniles. These results suggest that the prefrontal to dorsal raphe pathway is a critical anatomical and functional substrate of the development and manifestation of behavioral control.

## Introduction

The emergence of behavioral control, including attention, patience, cognitive flexibility, and behavioral persistence, occurs during critical periods of postnatal development. In these phases, environment and experience contribute to the maturation of higher cognitive functions (Larsen and Luna, 2018; Mischel et al., 1989; Tooley et al., 2021), which sets the foundations of future social and cognitive abilities during adulthood (Casey et al., 2011; Moffitt et al., 2011).

Ethologically, the development of behavioral control is critical for selective fitness and, thus, survival. For instance, in the natural environment, food resources are often sparsely distributed and depleted with consumption. Therefore, the well-known tradeoff between exploiting a depleting resource and exploring in search of alternatives is crucial to reach an optimal foraging strategy and obtain the maximum amount of resources with minimal waste of physical effort. Therefore, a forager in a possibly depleted patch of food faces an important dilemma--to stay or to leave--that calls for a careful balancing between persistence and flexibility (Charnov, 1976; Lottem et al., 2018; Morris and Davidson, 2000; Vertechi et al., 2020).

From a neural perspective, cognitive development correlates with large-scale synaptic and structural changes (Durston and Casey, 2006; Zuo et al., 2010, 2017) that are considered to underlie the emergence of increasing cognitive control over innate impulsive behavioral tendencies (Alexander-Bloch et al., 2013; Fair et al., 2009; Luna et al., 2001). A variety of evidence links the medial prefrontal cortex (mPFC) to the expression of behavioral control in a wide range of mammal species. For instance, humans and macaques with prefrontal cortical damage display deficits in behavioral flexibility, decision making, and emotional processing (Izquierdo et al., 2017; Rudebeck et al., 2013; Roberts et al., 1998), as well as a notable increase in impulsive behavior (Berlin, 2004; Dalley and Robbins, 2017; Fellows, 2006; Itami and Uno, 2002). In line with this, local pharmacological inhibition of mPFC significantly limits rats’ ability to wait for a delayed reward (Murakami et al., 2017; Narayanan et al., 2006).

Crucially, the mPFC undergoes intense postnatal maturation from childhood to adulthood, particularly during adolescence (Chini & Hanganu-Opatz., 2021), which in humans spans from years ∼10-18 of life and in mice from weeks ∼3-8 of life, and is a period of intense somatic maturation, including sexual development (Bell, 2018).

The maturation of the mPFC includes structural and functional modifications that overlap in time with the development of patience and behavioral inhibition during childhood and adolescence (Chini and Hanganu-Opatz, 2021; Sakurai and Gamo, 2019). Although it has been long hypothesized that the neural changes occurring in the mPFC during development are central to the emergence of behavioral control (e.g. Durston and Casey, 2006; Sowell et al., 1999), the specific plastic arrangements underlying behavioral control development remain poorly understood.

A number of studies have focused on the local changes of mPFC circuits, such as the changes in cortical thickness caused by cellular structural plasticity and synaptic pruning that are characteristic of early postnatal developmental phases in both humans and mice (Nagy et al., 2004; Alexander-Bloch, 2013; Ueda et al., 2015; Kolk & Rakic 2021), as a putative locus underlying cognitive development. More recently, studies in rodents have shed light on the development of long-range top-down mPFC extracortical connections as the putative origin of certain aspects of cognitive development (Klune et al., 2021). In particular, the development of mPFC afferents to the amygdala may shape the response to threats across different stages of development (Arruda-Carvalho et al., 2017; Dincheva et al., 2015; Gee et al., 2016), and the development of mPFC input onto the dorsal raphe nucleus (DRN) shapes the response to stress (Soiza-Reilly et al., 2019).

A growing body of evidence supports that 5HT neuron activity in the DRN is related to increases in the ability to wait for rewards (Fonseca et al., 2015; Lottem et al., 2018; Miyazaki et al., 2011, 2018, 2014). This reflects a prolongation of the willingness of animals to engage in an active behavior such as foraging, rather than promotion of passivity (Lottem et al. 2018), and can also involve active overcoming of adverse situations (Nishitani et al., 2019; Ohmura et al., 2020, Warden et al., 2012).

The mPFC sends a dense glutamatergic projection to the DRN at the adult stage (Pollak Dorocic et al., 2014; Weissbourd et al., 2014; Zhou et al., 2017), which can bidirectionally modulate the activity of 5HT DRN neurons through monosynaptic excitation or disynaptic feedforward inhibition through local interneurons (Challis et al., 2014; Geddes et al., 2016; Maier, 2015; Warden et al., 2012). Selective optogenetic activation of the mPFC inputs to the DRN elicits active behavioral responses in a challenging context (Warden et al., 2012), and perturbations in the development of this pathway lead to maladaptive anxiety levels (Soiza-Reilley et al., 2019).

Given the reciprocal connectivity between the mPFC and DRN (Puig and Gulledge, 2011) and that both areas causally modulate animals’ ability to wait for delayed rewards (Ciaramelli et al., 2021; Fonseca et al., 2015; Miyazaki et al., 2011, 2018; Murakami et al., 2017; Schweighofer et al., 2008), it seems plausible that the maturation of mPFC input to the DRN over development could underlie the emergence of behavioral persistence in mice.

Therefore, we sought to characterize the development of cortical innervation onto the DRN and its functional consequences in the context of behavioral persistence. In contrast to previous studies in which adult behavioral readouts were assessed after developmental perturbations (Bitzenhofer et al., 2021; Soiza-Reilly et al., 2019), we undertook a longitudinal study, characterizing behavior, synaptic physiology and anatomy, in parallel, from adolescence to adulthood.

The study of cognitive development in mice is challenged by the fact that most tasks designed to assess cognition require several weeks or even months of training (Pittaras et al., 2020; Sanchez-Roige et al., 2012; The International Brain Laboratory et al., 2021; Winstanley and Floresco, 2016). Here, we took advantage of the innate foraging ability to test naive mice in a two-port probabilistic foraging task which measured their persistence in exploiting a given foraging port before exploring the other. This task was used previously in adult mice to test their ability to infer the hidden state of the foraging site, a behavior that took several days to establish (Vertechi et al., 2020), Here, instead, we took advantage of the fact that even naive mice performed the task competently, although sub-optimally, from the first day of exposure. We found that all mice were able to perform the movements required to obtain water within minutes of entering the behavioral box, including nose-poking in the ports, licking at the water spout, and alternating between ports, suggesting that this task likely taps into a largely innate behavioral repertoire.

We show that the behavior of mice grows more persistent from juvenile (3-4 weeks old) to adulthood (7-8 weeks), that these behavioral changes mirror the time course of maturation of the mPFC to DRN projection, and that ablation of this projection in adult mice recapitulates the juvenile behavioral phenotype. These results suggest that the development of top-down PFC-DRN afferents is critical to the emergence of cognitive control over behavioral impulsivity that characterizes adulthood.

## Results

### Cortical top-down input over the dorsal raphe matures in the transition between adolescence and adulthood in mice

First, to characterize the development of neocortical projections to the DRN, we focused on the afferents of layer V neurons, which are the primary origin of these projections (Pollak Dorocic, 2014). Using a mouse line expressing channelrhodopsin-2 (ChR2) in a large fraction neocortical layer V neurons (Rbp4-Cre/ChR2-loxP) <u>(Leone et al., 2015)</u>, we performed ChR2-assisted circuit mapping (sCRACM) <u>(Petreanu et al., 2009)</u> of cortical afferents in brain slices containing the DRN obtained from mice between postnatal weeks 3 to 12 (Fig 1A). Taking advantage of the fact that ChR2-expressing axons are excitable even when excised from their parent somata, we evoked firing of presynaptic ChR2-expressing cortical axons innervating the DRN while recording the electrophysiological responses of postsynaptic DRN neurons. We assessed the fraction of recorded DRN neurons receiving cortical excitatory synaptic input (connection probability, Pcon) and the strength of this connection (amplitude of the evoked synaptic response) at different developmental time points.

**Figure 1.**
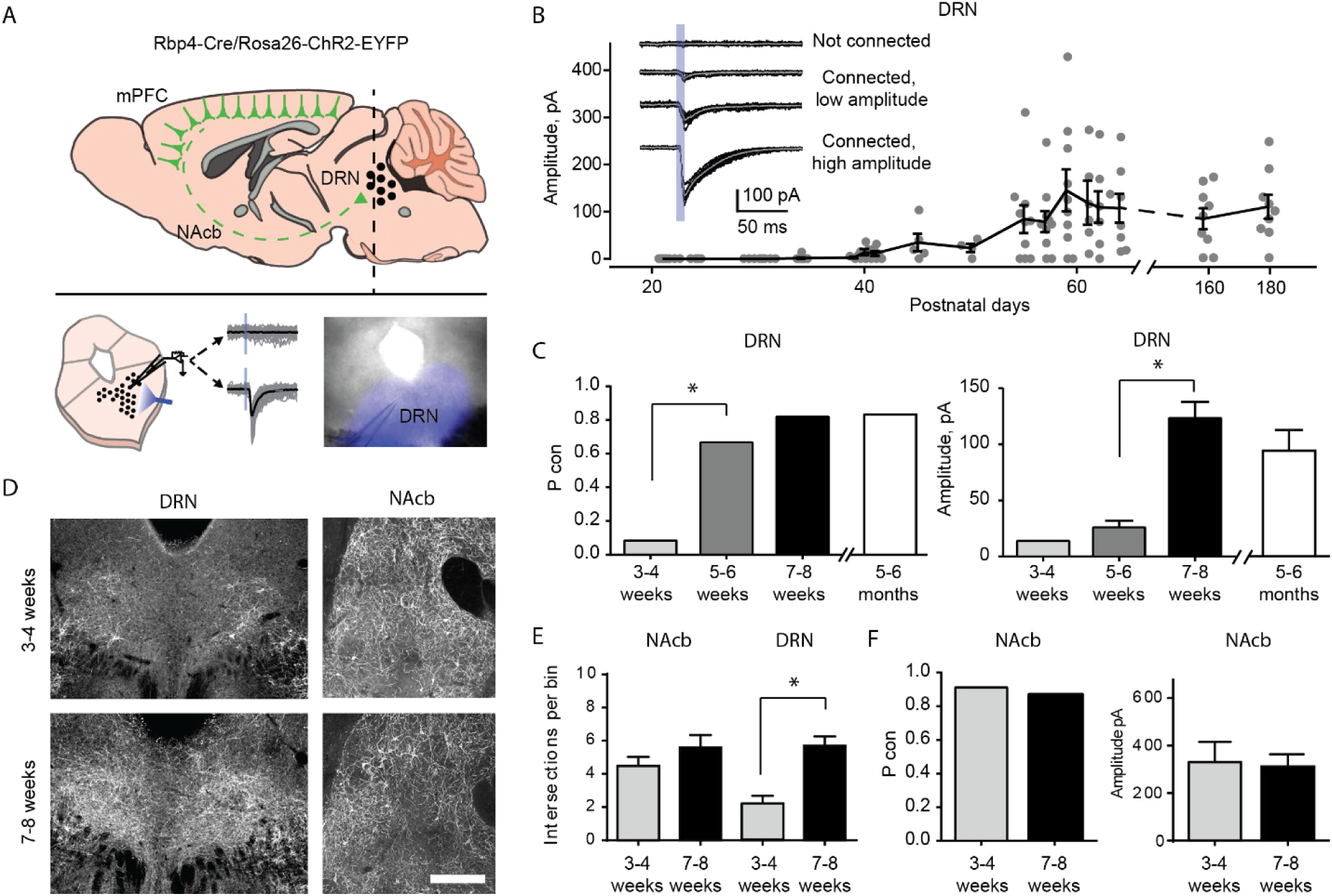
Top-down cortico-raphe connections develop over adolescence in mice. (A) Schematic representation of a sagittal view of an Rbp4-ChR2 mouse brain illustrating top-down cortico-raphe afferents. Coronal slices containing the DRN were obtained ex vivo, and whole-cell recordings of DRN neurons were performed to assess cortical connectivity upon light stimulation. (B) Optogenetically evoked EPSCs were recorded in DRN neurons contacted by ChR2 expressing cortical axons (122 neurons, 20 Rbp4-ChR2 mice). The current amplitude of cortico-raphe connections is plotted as a function of postnatal age in mice. (C) Pooled connection probability (Connected cells/ total cells) and averaged connection amplitude of cortico-DRN afferents at four different developmental points: early juvenile (3-4 weeks), late juvenile (5-6 weeks), early adult (7-8 weeks), late adult (5-6 months). (D) Example images illustrate an increased cortico-DRN innervation in adult mice compared to juveniles, while the cortico-accumbens innervation remains constant over the same time period. Scale bar = 400 microns. (E) Number of axonal intersections quantified in the DRN and nucleus accumbens of juvenile and adult mice. (F) Pooled connection probability and averaged connection amplitude of cortico-accumbens afferents in early juvenile and early adult mice. * indicates p<0.05.

We found a dramatic increase in the connection probability and amplitude of cortico-raphe input between weeks 3 and 8 (Fig. 1B-C). Between the 3-4 weeks (juvenile mice), the probability of DRN neurons receiving cortical input was equal to 0.07. This probability increased significantly to 0.66 (Pcon 3-4 weeks vs. Pcon 5-6 weeks, Chi-Square test Χ^2^ (1, N = 53 neurons) = 24.1, p = 0.00001) between weeks 5 and 6, reaching a peak connection probability of 0.82 between weeks 7 and 8 (Fig. 2C). Between 5-6 and 7-8 weeks (i.e. late juvenile to adult mice), the amplitude of the optogenetically evoked currents increased from 27.3 ± 6.2 pA to 128 ± 15.7 pA (mean ± SEM, two-tailed t-test, t(57) = 4.03, p = 0.002). To test whether there is a further development of this pathway in the later stages of development, we recorded slices from 12-week old mice. We observed no further increase in either the connection probability (Pcon 7-8 weeks = 0.82 vs.Pcon 5-6 months = 0.80, Chi-Square test Χ^2^ (1, N = 70 neurons) = 0.03, p = 0.84) or the input magnitude (7-8 weeks old = 126 ± 15 pA vs. 5-6 months old = 113 ± 14.1 pA, two-tailed t-test, t(60) = 1.27, p = 0.21, Fig. 2B-C). Altogether, these results suggest that the cortico-raphe pathway gradually matures between weeks 3 and 8 and then plateaus. Importantly, the location of the recorded DRN neurons was comparable between juvenile and adult mice (Fig. S1) and thus, the connectivity changes observed across development do not reflect a biased sampling of differentially innervated sub-regions of the DRN. Furthermore, we observed a comparable input resistance (3-4 weeks: median = 444 MΩ, 95% CI = [370, 676], 5-6 weeks: median = 612 MΩ, 95% CI = [402, 925], 7-8 weeks: median = 731 MΩ, 95% CI = [519, 943], 5-6 months: median = 532 MΩ, 95% CI = [385, 664], Kruskal-Wallis H(3) = 6.06, p = 0.11) and input capacitance (3-4 weeks: median = 20.7 pF, 95% CI = [17.4, 25.9], 5-6 weeks: median =22.8 pF, 95% CI = [15.9, 24.8], 7-8 weeks: median = 20.8 pF, 95% CI = [18.3, 28.5], 5-6 months: median =23.5 pF, 95% CI = [14.1, 44.7], Kruskal-Wallis H(3) = 0.81, p = 0.84) in DRN neurons over development (Fig. S2A,B), suggesting that changes at the level of the passive propagation of current through DRN neurons is not the underlying cause of the increase in connection probability and input magnitude observed over time.

**Figure 2.**
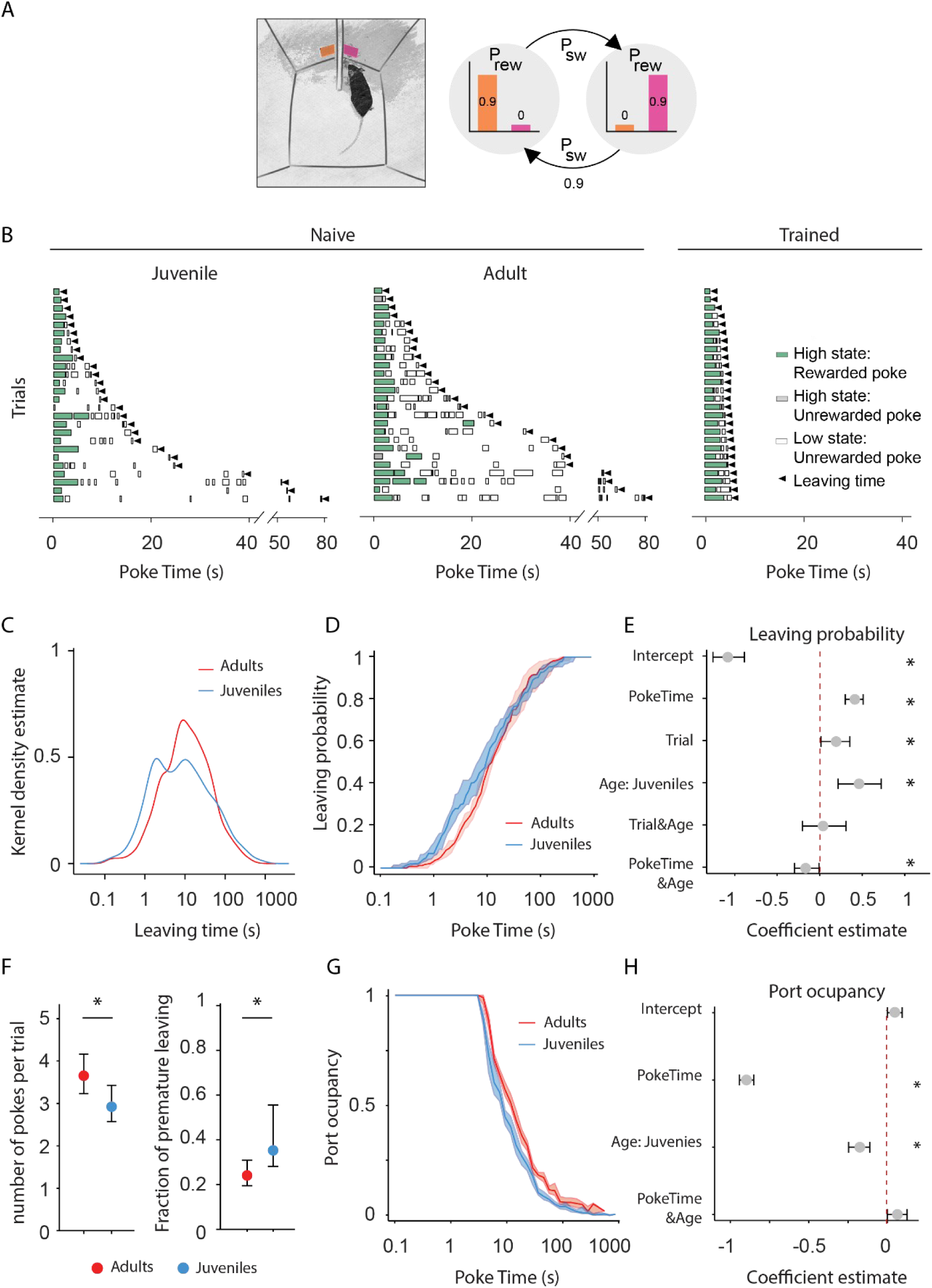
Adult mice persist longer than juveniles in exploiting a foraging patch. **(A)** Illustration of the rodent foraging task. Water-deprived mice seek rewards by probing two nose-ports. (B) Randomly selected examples of poking behavior throughout a naive juvenile, naive adult and trained adult behavioral session sorted by trial length. Pokes in the active state can be rewarded (in green) or not (in gray). Pokes in the inactive state are never rewarded (in white). After the state switches, the mice have to travel to the other side (left or right port, L and R) to obtain more water. Leaving time is illustrated with black triangles. (C) Distribution of the trial durations for naive juveniles and naive adults. (D).Cumulative distribution of the probability of leaving (median ± 95% CI across mice) as time elapses from the first poke in a trial for adults and juvenile animals. (E) Regression coefficients ± 95% CI resulting from a parametric bootstrap (n = 1000) of a mixed models logistic regression to explain the probability of leaving. * indicates predictors with a significant impact on the probability of leaving. (F) Median ± 95% CI fraction of the number of pokes per trial (left) and incorrect trials (right). Juvenile mice do a significantly lower amount of pokes per trial and a higher proportion of premature leaving. (G) Port occupancy as a function of trial time elapsed for juveniles and adults. (H) Regression coefficients ± 95% CI resulting from a parametric bootstrap (n = 1000) of a mixed models logistic regression to explain the port occupancy, as in F. All analyses in C-H computed by pooling the data from all sessions of juvenile (N = 21) or adult (N = 23) mice, yielding a total of 2875 trials (juveniles = 1347, adults = 1528) and 9596 pokes (juveniles = 3908, adults = 5688).

In these experiments, the onset of ChR2 expression is dictated by the Cre recombinase expression under the control of the native Rbp4 promoter over development. Therefore, if in the juvenile cortex there were fewer neurons expressing Rbp4 or the onset of expression was near our recording time point, this could affect the net amount of ChR2-expressing top-down cortical axons and/or their net excitability. To control that our findings reflect a development process and not a genetic artifact caused by the temporal dynamics of Rbp4 expression, we performed two additional control experiments in one of the main cortical origins of afferents onto the DRN, the mPFC (Weissbourd et al., 2014, Zhou et al., 2017).

First, we compared the density of neurons in the mPFC expressing a fluorescent reporter (tdTomato) under the control of the Rbp4 promoter in juvenile and adult mice. The same density of tdTomato expressing somas was detected in the mPFC of juvenile and adult Rbp4-Cre\ tdTomato-loxP mice (Juveniles: median = 4.59 somas per 0.01 mm^2^, 95% CI = [4.59, 6.18], vs. Adults: median = 5.22 somas per 0.01 mm^2^, 95% CI = [4.35, 6.06], Mann-Whitney U test (N_Juveniles_ = 3,N_Adults_ = 4) = 6, p = 0.99, Fig. S2C), indicating that a comparable number of neurons underwent a Cre-dependent recombination of the tdTomato fluorescent reporter under the control of the Rbp4 promoter at both developmental time points. Second, we compared the light-evoked somatic current produced in layer V neurons expressing ChR2 under the Rbp4 promoter in juvenile and adult mice. In agreement with the previous control, layer V neurons in the mPFC expressing ChR2 under the Rbp4 promoter produced the same amount of photocurrent upon light stimulation in juvenile and adult mice (Repeated Measures ANOVA, F(1, 11) = 0.138, p = 0.71 for age factor, Fig. S2D). These experiments show that juvenile and adult mice have similar densities of cortical layer V projection neurons that could give rise to DRN afferents and that these neurons express similar amounts of ChR2 and thus, if present, projections should be equally detectable by optogenetic circuit mapping across ages. Altogether, this evidence suggests that the maturation of cortico-raphe projections we report is caused by a developmental process and is not explained by experimental artifacts.

To further understand the mechanisms underlying the cortico-raphe input strengthening observed over development, we investigated whether the changes of connection probability and input amplitude we observed were accompanied by differences in the density of cortical axonal innervation over the DRN. Indeed, we observed a significantly higher density of Rbp4 positive axons around the DRN in adults compared to juveniles (2.2 ± 0.47 vs. 5.7 ± 0.56 axons per bin in juvenile vs.adult mice, two-tailed t-test (N_Juveniles_ = 4,N_Adults_ = 7), t(9) = 4.15, p = 0.002, Fig. 1D-E). This observation supports the idea that the increase in physiological strength we observed reflects in part the growth of new connections between the neocortex and DRN.

To assess whether the development of cortico-raphe projections is specific to raphe projecting cortical afferents or it reflects a more general maturation of corticofugal projections over adolescence in mice, we mapped the anatomical and synaptic development of cortico-accumbens projections, which are mainly originated in the PFC (Phillipson & Griffiths 1985, Li et al., 2018) and which functional connectivity has been previously assessed in juvenile rodents (Gorelova and Yang, 1996). In contrast to cortico-raphe afferents, cortico-accumbens projections did not undergo any significant structural change over the same developmental period (4.5 ± 0.54 vs. 5.8 ± 0.74 axons per bin in juvenile vs. adult mice, two-tailed t-test (N_Juveniles_ = 3, N_Adults_ = 7), t(8) = 1.09, p = 0.40, Fig. 1D-E). Consistent with the anatomy, the ChR2-assisted mapping of cortico-accumbens connections in juvenile and adult Rbp4-Chr2 mice revealed no change in either the connection probability (Pcon 3-4 weeks = 0.90 vs.Pcon 7-8 weeks = 0.87, Chi-Square test Χ^2^ (1, N = 19 neurons) = 0.03, p = 0.81) or the input amplitude in the transition from juveniles to adults (two-tailed t-test, t(15) = 0.15, p = 0.88, Fig. 1F). Altogether, these observations reveal the structural and synaptic development of a subpopulation of cortical afferents targeting the DRN during the transition from juvenile to adult in mice that does not reflect a generalized development of corticofugal projections.

### Baseline persistence correlates with the maturation of cortico-raphe input in the transition between adolescence and adulthood in mice

To investigate the development of behavioral persistence in mice, we employed a previously published self-paced foraging task (Vertechi et al., 2020). The setup consists of a box with two nose-ports separated by a barrier (Fig. 2A). Each nose-port constitutes a foraging site that mice can actively probe in order to receive water rewards. Only one foraging site is active at a time, delivering reward with a certain probability (Prwd = 90%) when probed. Each try in the active site can also cause a switch of the active site’s location with a certain probability (Psw = 90%) (Fig. 2B). After a state switch, mice have to travel to the other port to obtain more reward, bearing a time cost to travel. While this task was previously studied after multiple days of training, after which adult mice use an inference-based strategy, early in learning, they show a value-based strategy, staying longer when they receive more rewards (Vertechi et al., 2020). This behavior was useful for the purpose of assessing cognitive development, as it requires persistence in poking at the port despite reward failures.

To measure how developmental changes affect mice’s persistence, we compared the behavior of juvenile (weeks 3-4) and adult mice (weeks 7-8) on their first exposure to the apparatus and task. Presumably due to the novelty of the apparatus, mice tended to interleave poking in the port with investigating the apparatus, often taking long pauses in between pokes at the same port. This resulted in a less regular poking structure than experienced mice (Fig. 2B). However, both adults and juveniles succeeded in performing the required actions of poking and traveling between ports, receiving substantial rewards over the course of the session and there were no gross differences in the behavior of juvenile and adult mice (Fig. 2B).

In order to compare behavioral persistence across development we first assessed the animals’ leaving time, measured as the overall time spent investigating one port before visiting the other (time elapsed from the first to the last poke in a port, as illustrated in Fig. 2B). There was no difference in the mean or median leaving time for juveniles vs. adults. However, inspection of the distributions of leaving times showed an extremely heavy tail of long ’site visits’ (Fig. 2C) Assuming that very long leaving times reflect not continued foraging episodes but behavioral ’lapses’ due to exploration or other distractions, we applied an arbitrary cutoff of 60s to both distributions and compared the medians of the resulting truncated distributions. The comparison revealed that juveniles had significantly shorter median leaving times (Adults: median = 0.96, 95% CI = [0.05, 0.06], Juveniles: median = 0.78, 95% CI = [0.15, 0.06]; Mann Whitney U test (N_Adults_ = 21, N_Juveniles_ = 23) = 366, p = 0.0029, effect size = 0.76), indicative of reduced persistence. This could also be seen in a comparison of cumulative distributions (Fig. 2D), which shows a leftward shift in juveniles for trials around 1-10 s, a time scale which is the typical duration of trained animals’ trials.

To formalise this analysis without the use of arbitrary cutoffs, we performed a logistic regression for probability of leaving the patch as a function of the Time within the trial, the Age of animal (juvenile vs. adult) and elapsed trials within the session (Trial):

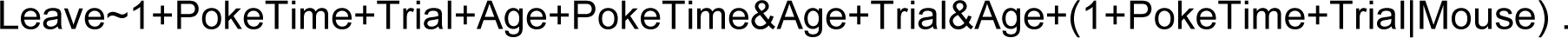

The individual variability was accounted for through generalized linear mixed models with random intercept and slopes for each mouse (see methods for the implementation). A factor was considered to significantly affect the decision to leave if the value of its estimated coefficient plus 95% confidence interval (1000 parametric bootstrap analysis, see methods) did not cross 0. This analysis showed a significant effect of Age, with juveniles more likely to leave than adults (Fig. 2E). We confirmed that the animals’ Age group significantly contributes to the ability to explain the probability of leaving (likelihood ratio test on Leave∼1+PokeTime+Trial+Age+PokeTime&Age+Trial&Age+(1+PokeTime+Trial|MouseID) versus Leave∼1+PokeTime+Trial+(1+PokeTime+Trial|MouseID): Χ^2^_(3)_ = 19.27, p = 2e^-4^). The probability of leaving increased as a function of Trial, indicating that animals become less persistent over the course of the session (Fig. S3A). Including the Trial factor also improved the model prediction (likelihood ratio test on Leave∼1+PokeTime+Trial+Age+PokeTime&Age+Trial&Age+(1+PokeTime+Trial|MouseID) versus Leave∼1+PokeTime+Age+PokeTime&Age+(1+PokeTime|MouseID): Χ^2^_(5)_ = 128.21, p < 1e^-25^). This effect could suggest a drop in motivation due to the water drunk during the session. However there was no significant interaction between Trial and Age factors, indicating that differences in satiety, fatigue or learning accumulated throughout the session do not underlie the change in persistence between juveniles and adults.

The use of leaving time as a measure of persistence does not take into account the nature of the behavior before leaving. A difference in persistence could also manifest as differences in the fraction of time spent actively poking. Indeed, adults make more pokes per trial than did juveniles (Fig. 2F; Mann-Whitney U test (N_Adults_ = 21, N_Juveniles_ = 23) = 354.5, p = 0.008, effect size = 0.73). Next, to assess port occupancy, we quantified the cumulative time spent with the snout in the port divided by the overall time elapsed from the trial beginning (Fig. 2G). This analysis also showed a clear difference between juvenile and adult mice, with adults’ port occupancy extending long into the trial. Consistent with the leaving time regression results, port occupancy decreased with time into a trial. and juveniles’ poke occupancy decreased significantly faster than the adults’, as indicated by both a significant effect of the Age variable (Fig. 2H) and significant improvement in regression with the Age factor: likelihood ratio test on Occupancy∼1+PokeTime+Age+Age&PokeTime+(1+PokeTime|MouseI vsOccupancy∼1+PokeTime+(1+PokeTime|MouseID), Χ^2^_(2)_ = 20.10, p < 1e^-4^).

These results indicate a difference in persistence of foraging behavior in juvenile vs. adult mice. We also found that the difference in behaviour between juveniles and adults had an impact on performance. Compared to adults, juveniles performed a larger proportion of incorrect trials, i.e. leaving before the reward side had switched (Fig. 2F; Adults: median = 24% incorrect, 95% CI = [24%, 31%], Juveniles: median = 36% incorrect, 95% CI = [28%, 55%]; Mann-Whitney U test (N_Adults_ = 21, N_Juveniles_ = 23) = 137.5, p = 0.015, effect size = 0.72). This would presumably result in a reduction in foraging efficiency, but, since both juvenile and adult naive animals’ exhibited substantial periods of non-foraging exploratory behavior, we did not attempt to further quantify this.

Finally, to assess whether the developmental changes are consistent in male and female mice we tested the variable sex in the analysis of the previous cohort of mice. We found no significant effect of Sex on the probability of leaving either alone or in interaction with animals’ age (Fig. S3C), and including Sex as a predictor had no significant improvement in the model’s ability to explain the decision to leave (Fig. S3D, likelihood ratio test on Leave∼1+PokeTime+Trial+Age+Sex+Age&Sex+(1+PokeTime+Trial|MouseID) versus Leave∼1+PokeTime+Trial+Age+(1+PokeTime+Trial|MouseID): Χ^2^_(2)_ = 3.46, p = 0.18). These results suggest that the maturation of persistence occurs at a similar rate in male and female mice.

### mPFC-DRN pathway ablation in adult mice recapitulates juvenile behavioral features

The above results establish a correlation between the development of the descending cortical input to the DRN and the emergence of behavioral persistence. To more directly causally link the development of cortico-raphe afferents to the increase in persistence observed in the probabilistic foraging task, we next ablated the cortico-raphe pathway in adult mice and assessed the impact on behavioral persistence.

To ablate cortico-raphe afferents, we used an engineered version of Caspase3 (taCasp3-TEVp) that is able to trigger apoptosis bypassing cellular regulation upon activation by the TEV protease, which is coexpressed in the same construct (Yang et al., 2013). We packaged a Cre dependent taCasp3-TEVp construct (or the reporter tdTomato as a control) in a retrogradely travelling AAV vector (rAAV), that we locally delivered in the DRN of Rbp4-Cre mice. This approach resulted in the fluorescent tagging of cortico-raphe layer V projecting neurons in control mice (tdTomato mice) and in the ablation of the same corticofugal pathway in taCasp3-TEVp injected mice (Caspase mice) (Fig. 3A).

**Figure 3.**
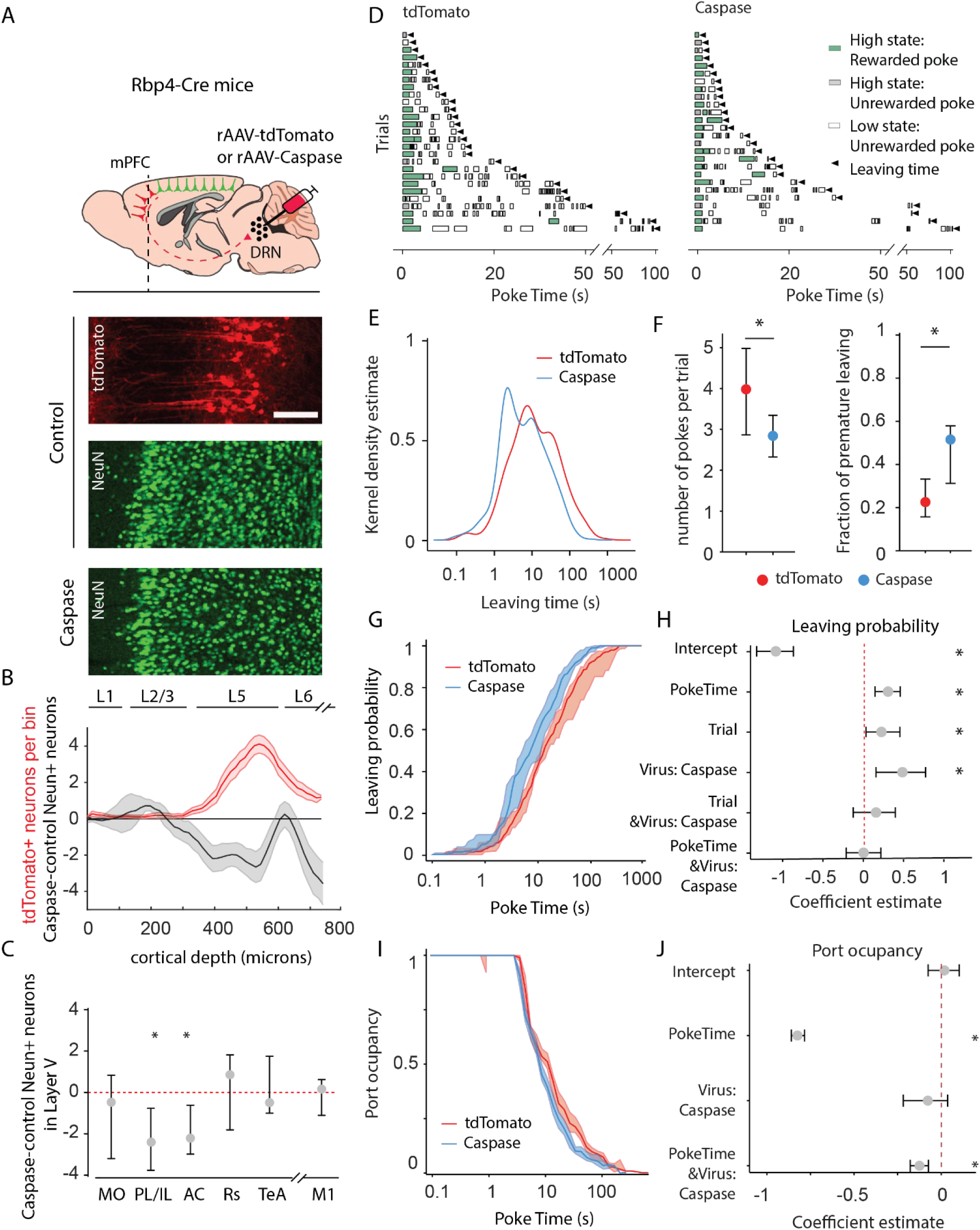
Animals lacking cortico-raphe projections show less behavioral persistence in exploiting a foraging patch. (A) Schematic representation of the ablation strategy used for behavioral assessment. Retrogradely transporting AAV vectors expressing either the fluorescent reporter tdTomato (rAAV-tdTomato) or the intrinsically active apoptosis triggering Caspase3 (rAAV-Caspase, Caspase group were locally delivered in the DRN of Rbp4-Cre mice. In the cortical areas containing tdTomato expressing neurons in control animals (in example picture, PL/IL cortex) the density of neurons was quantified with an immunohistochemistry protocol against the pan-neuronal marker NeuN and compared to the neuronal densities obtained in the same cortical areas of ablated mice (scale bar = 200 microns). (B) Distribution of the neuronal density difference between ablated mice and the mean density of control mice per cortical depth bin in the PL/IL cortex (black shaded error plot). The neuronal density loss observed in ablated mice when compared to control NeuN densities matches the cortical depth in which tdTomato neurons are located (red shaded area). Shaded error plots represent mean ± SEM. (C) Summary of caspase-control NeuN density per brain area (MO: median = -0.46, 95% CI = [-2.81, 1.06], PL/IL: median = -2.3, 95% CI = [-4.01, -1.44], AC: median = -2.26, 95% CI = [-3.59, -0.17], Rs: median = 0.75, 95% CI = [-1.80, 1.81], TeA: median = -0.63, 95% CI = [-1.01, 1.76], and M1: median = 0.14, 95% CI = [-1.81, 0.73]). (D) Randomly selected examples of poking behavior for a tdTomato and caspase behavioral session sorted by trial length. Pokes in the active state can be rewarded (in green) or not (in gray). Pokes in the inactive state are never rewarded (in white). Leaving time is illustrated with black triangles. (E) Distribution of the trial durations for tdTomato and caspase mice. (F) Median ± 95% CI fraction of the number of pokes per trial (left) and incorrect trials (right). Caspase mice do a significantly lower amount of pokes per trial and a higher proportion of premature leaving. (G) Cumulative distribution of the probability of leaving as a function of trial time elapsed (median ± 95% CI across mice) for tdTomato and Caspase animals. (H) Regression coefficients ± 95% CI resulting from a parametric bootstrap (n = 1000) of a mixed models logistic regression to explain the probability of leaving. (I) Port occupancy as a function of trial time elapsed for tdTomato and Caspase. (J) Regression coefficients ± 95% CI resulting from a parametric bootstrap (n = 1000) of a mixed models logistic regression to explain the port occupancy. All analyses in B-G computed by pooling the data from the histology and the first session of Caspase (N = 7) or tdTomato (N = 8) mice, yielding a total of 1464 trials (Caspase = 939, tdTomato = 525) and 4742 pokes (Caspase = 2555, tdTomato = 2187).

The prelimbic/infralimbic (PL/IL) and anterior cingulate (AC) cortices, which constitute the mPFC, were the areas with the highest density of DRN-projecting tdTomato+ somas in control animals and consistently more extensive neuron density loss in caspase injected mice, quantified using the pan-neuronal marker NeuN (Fig. 3A-C, Fig. S4, control vs. caspase, two-sample Kolmogorov-Smirnoff Test = 0.028, p = 0.002 for PL/IL and D = 0.024, p = 0.01 for AC). We also found tdTomato+ somata in the medial orbitofrontal cortex (MO) of the control group; however, this projection was weaker in terms of tdTomato+ labelled neurons and, consistently, the difference in layer V NeuN densities between control and caspase mice was not significant (Figs. 3C, S4, D = 0.017, p = 0.08).

Apart from the mPFC, sparse labeling of tdTom+ neurons was found in more posterior levels of the neocortex, namely in the retrosplenial cortex (RS) and in the temporal association cortex (TeA) (Fig. S4). Nonetheless, tdTom+ neurons in the RS and TeA were found in only 5 and 3 out of 8 control animals, respectively. Consistently, the reduction in NeuN layer V neuronal density in these two areas was minimal and non-significant compared to controls (Fig. 3C, D = 0.034, p = 0.12 for RS and D = 0.025, p = 0.19 for TeA). In addition, no differences in NeuN density were observed between caspase injected animals and controls in a control area not showing tdTomato expressing somas and therefore not projecting to the DRN (M1, Fig. 3C, D = 0.019, p = 0.15). These observations suggest that our ablation approach significantly affected mPFC-DRN projecting neurons, particularly from PL/IL and AC cortices.

When investigating the distribution of tdTomato expressing somas, we observed weak collateral projections of the cortical subpopulation projecting to the DRN in the lateral septum, lateral hypothalamic nucleus, the ventral tegmental area and the anterior periaqueductal gray; medium collateral axonal density in anterior subcortical olfactory nuclei (anterior dorsal endopiriform, anterior olfactory nucleus, dorsal taenia tecta and islands of Calleja) and the substantia nigra; and heavy collateralization in the dorsomedial striatum (Fig. S5).

We then assessed the impact of ablation of the mPFC-DRN projection on behavioral persistence using the foraging paradigm. We observed a similar pattern of differences between caspase and tdTomato mice as between juveniles and adults (Fig. 3D-F). Caspase animals showed an increase in the probability of leaving the port, seen as a leftward shift in the cumulative distribution of leaving times (Fig. 3G). We applied logistic regression analysis to identify which factors significantly affect the probability of leaving after each poke. Both PokeTime and Trial significantly influenced the probability of leaving (Fig. 3H). Crucially, we found that animals lacking cortico-raphe projections are significantly more likely to leave the patch earlier than control animals (Fig. 3H). As for the comparison of juvenile and adult mice, this effect did not interact with PokeTime (Fig. 3H). Including the Virus group factor significantly improved the explanatory power of the model (likelihood ratio test on Leave∼1+PokeTime+Trial+Virus+PokeTime&Virus+Trial&Virus+(1+PokeTime+Trial|MouseID) versus Leave∼1+PokeTime+Trial+(1+PokeTime+Trial|MouseID): Χ^2^_(2)_ = 10.84, p = 0.012). Interestingly the reduced persistence of Caspase animals does not scale with the elapsed time as for the juveniles (lack of interaction effect PokeTime&Virus, Fig. 3H).

We also characterized differences in poking behavior. As with juvenile vs. adult mice, caspase mice made significantly fewer pokes per trial than adults (Fig. 3F; Mann-Whitney U test (N Caspase = 7, N tdTomato = 8) = 49, p = 0.01, effect size = 0.82) and showed a shift toward shorter port occupancy (Fig. 3I). A regression analysis showed that the viral intervention impacted significantly poke occupancy (likelihood ratio test on Occupancy∼1+PokeTime+Virus+Virus&PokeTime+(1+PokeTime|MouseID) vs Occupancy∼1+PokeTime+(1+PokeTime|MouseID), X2(2) = 12.69, p = 0.018), the regression analysis showed that this was mainly due to a progressive reduction during the trial rather than a subtractive effect (significant PokeTime & Virus: Caspase, not Virus: Caspase alone, Fig. 3J).

Finally, we assessed whether ablating the prefrontal-DRN pathway changed the performance on the task. Compared to tdTomato mice, caspase mice committed a significantly larger proportion of errors, prematurely leaving the site before the active port switches (Fig. 3F; tdTomato: median = 22% incorrect, 95% CI = [16%, 33%], caspase: median = 50% incorrect, 95% CI = [32%, 57%]; Mann Whitney U test (N_tdTomato_ = 8, N_Caspase_ = 7) = 137.5, p = 0.01).

Together these results indicate that ablating the mPFC-DRN pathway recapitulates the key behavioral features characteristic of juvenile mice in the same foraging task, indicating that the mature mPFC-DRN projection is necessary for the behavioral persistence displayed by adult mice and suggesting that this pathway is likely to contribute to the development of behavioral persistence in mice.

## Discussion

In the present study, we described how the postnatal maturation of the mPFC projection to the DRN during adolescence is linked to the performance of a probabilistic foraging task. Over the same period of development, the mPFC to DRN projection underwent a dramatic increase in potency and mice developed an increase in persistence in foraging behavior. Ablation of the mPFC-DRN pathway in adult mice recapitulated the features observed in the behavior of juvenile mice, supporting a causal relationship between the mPFC-DRN projection and behavioral persistence.

In a wide variety of species, including mice, adolescence corresponds to the emancipation from the parents (Spear, 2000), a period in which individuals need to develop or refine skills to become independent. This ethological scenario may explain the evolutionary selection of juvenile behavioral traits (Sercombe, 2014; Spear, 2000), such as increased impulsivity or high risk taking behavior (Laviola et al., 2003; Sercombe et al., 2014). However, the abnormal development of cognitive control over the intrinsic behavioral tendencies of juveniles may underlie aspects of the etiopathology of impulsive and addictive disorders in adult humans (Reiter et al., 2016, Wong et al., 2006). In line with pre-adolescent humans’ lack of delay gratification ability (Mischel et al., 1989), and with studies assessing impulsive behavior in mice over development (Sasamori et al., 2018), we found that mice of 3-4 weeks of age tend to be less persistent than 7-8 weeks old mice in a probabilistic foraging task. This led to a negative impact on foraging efficiency, with more premature site-leaving decisions.

From a neural perspective, maturational changes of prefrontal cortical areas, including the mPFC, have been previously linked to the emergence of cognitive skills during development in primates and humans (Luna et al., 2015, Nagy et al., 2004, Velenova et al., 2008). Such changes result in an increased top-down behavioral control and increased functional connectivity with cortical and subcortical targets (Hwang et al., 2010) in the transition between childhood to adulthood. However, the specific contribution of long-range top-down mPFC circuits and the cellular mechanisms underlying its development had not been previously investigated.

Here, using optogenetic-assisted circuit mapping we characterized the structural and functional development of cortico-raphe projections that take place over adolescence in mice. A recent report showed that a subpopulation of DRN-projecting mPFC neurons increases their axonal contacts over the DRN in an earlier phase of postnatal development (weeks 1-2) (Soiza-Reilly et al., 2019). We found a mPFC-DRN connection strength at 5-6 weeks postnatal similar to that reported by Soiza-Reilly and colleagues at a similar developmental time point (4-5 weeks of age). In addition, we found a connection strength at 7-8 weeks of age similar to those reported in adult rodents elsewhere (Zhou et al., 2017, Geddes et al., 2016). Thus, our findings are consistent with previous observations in the literature and suggest that the maturation of mPFC-DRN afferents starts early in postnatal development and undergoes an extended development period, plateauing only after reaching 7-8 weeks of age. Among the previous studies investigating the postnatal development of top-down afferents from the mPFC in rodents (Klune et al., 2021, Peixoto et al., 2016, Ferguson & Gao 2015), the latest mPFC afferent maturational process reported, the mPFC innervation over the basolateral amygdala, occurs up to week 4 (Arruda-Carvalho et al., 2017). Thus, to our knowledge, the mPFC-DRN pathway represents the latest top-down pathway from the cortex to develop.

Importantly, we found that the structural development of mPFC-DRN projections is causally linked to the maturation of behavioral persistence in adult mice. Using a genetically driven ablation approach (Yang et al., 2013), we selectively eliminated layer V cortical neurons projecting to DRN in adult mice. The procedure resulted in a behavioral phenotype that replicated key features of the juvenile foraging behavior. We observed a reduction in behavioral persistence, coupled with an increased fraction of errors. Furthermore, we localized the origin of these projections and quantified the local neuronal loss. The PL, IL, and AC cortices, areas that comprise the so called mPFC (Klune et al., 2021), suffered a significant loss with the procedure, highlighting their importance for displaying behavioral persistence necessary for reward exploitation in a foraging task.

Previous reports have shown that the pharmacological inactivation of the IL cortex reduces response persistence in a foraging task (Verharen et al., 2020). Moreover, lesions of the IL cortex improves performance on a reversal learning task (Ashwell and Ito, 2014), which resembles the increased behavioral flexibility observed in juvenile mice (Johnson and Wilbrecht, 2011). In addition, neurons in the PL cortex have been shown to track reward value and reflect impulsive choices (Sackett et al., 2017). More generally, the inactivation of PL/IL cortices using optogenetics leads to an increase in premature responses in a probabilistic reversal task (Nakayama et al., 2018), while the optogenetic activation of the PL/IL cortices increases food-seeking behavior while reducing impulsive actions (Warthen et al., 2016).

Furthermore, lesions in the AC impair behavioral inhibition producing an increase in premature actions in rodents (Muir et al., 1996, Hvoslef-Eide et al., 2018). More recently, it has been shown that the control of impulsive actions exerted by the AC requires intact signalling through Gi-protein in its layer-5 pyramidal neurons (Van der Veen et al., 2021). Altogether, there is considerable evidence linking the activity of the areas composing the mPFC (PL, IL and AC cortices) to the control of impulsive actions.

In addition, the optogenetic activation of DRN 5HT neurons, a major subcortical target of mPFC projections (Geddes et al., 2016; Zhou et al., 2017; Weissbourd et al., 2014, Pollak Dorocic et al., 2014), improves the performance of a delayed response task (Miyazaki et al., 2014, 2018; Fonseca et al., 2015) through an increase in active behavioral persistence (Lottem et al., 2018), which is the converse effect of the pharmacological silencing of the mPFC (Narayan and Laubach, 2006; Narayan et al., 2013; Murakami et al., 2017). Altogether, the emerging picture suggests that the individual activation of either mPFC or DRN converges into a behaviorally persistent phenotype. Consistent with this, the activation of mPFC-DRN top-down projections also has been shown to increase active persistence (Warden et al., 2012). However, previous studies have reported a net inhibitory effect of mPFC input onto 5-HT neurons in the DRN (Celada et al., 2001; Maier, 2015), particularly after prolonged trains of high-frequency stimulation (Srejic et al., 2015). This raises a question on the directionality with which mPFC input modulates DRN neuronal activity in the context of behavioral control. One possible mechanism would be a frequency dependency of the net effect, as found in thalamocortical connections (Crandall et al., 2015). In this scenario, given that the inhibitory interneurons in the DRN can track faster frequencies than 5-HT neurons (Jin et al., 2015) and that 5-HT neurons undergo 5-HT1a autoreceptor mediated inhibition upon dendritic NMDA receptor activation (De Kock et al., 2006), a prolonged activation of mPFC afferents to the DRN may, in turn, produce inhibition of 5-HT neurons. Nevertheless, other, less explored, patterns of cortical activity in different frequency ranges may tune 5-HT neuron subpopulations in different ways under more naturalistic patterns of activation and could be the focus of future research. An alternative mechanism for the bidirectional control of DRN activity by mPFC input would be synaptic plasticity, since it has been shown that activity dependent plasticity (Challis and Berton, 2015) and neuromodulators (Geddes et al., 2016) can bias the net excitatory or inhibitory effect that mPFC input exerts on DRN 5HT neurons.

In addition, while it’s well described that the PL/IL cortices produce a dense innervation over the DRN, the adjacent PR and IL cortices exert opposite effects on fear conditioning (Giustino & Maren, 2015) as well as on avoidance behaviors and behavioral inhibition (Capuzzo et al., 2020). This striking contraposition in their functional role leaves open the possibility of different circuit motifs on their DRN innervation that could explain a putative excitatory or inhibitory effect and that should also be the focus of future research.

The cortical subpopulation of DRN-projecting neurons manipulated in adult Rbp4-Cre mice in this study presented collateral projections that were particularly dense onto the dorsomedial striatum (Fig. S5), a pathway that has been shown relevant for foraging decisions (Bari et al., 2019). While it has been shown that the cortico-striatal pathway is fully developed after P14 using Rbp4-Cre mice (Peixoto et al., 2016) and therefore unlikely to underlie the developmental differences observed in this study, we cannot rule out an impact of the ablation of corticostriatal collaterals in the behavioral persistence decrease observed in Caspase treated mice.

The presence of parallel sub-systems in the DRN, with complementary projections either to the prefrontal cortex or to the amygdala and responsible for different behavioral responses has recently been reported (Ren et al., 2018). In our hands, mPFC-DRN descending neurons had very sparse collateralization to the amygdala (Fig. S5), while collaterals to the dorsal striatum or substantia nigra were abundant. This may suggest the presence of loops of preferential interconnectivity (mPFC→DRN/DRN→mPFC and mPFC→Amygdala/Amygdala→mPFC) as it has been shown for other cortical-subcortical loops (Young et al., 2021, Li et al., 2020), with different DRN subpopulations exerting specific neuromodulatory effects in either region (Ren et al., 2018).

To summarize, our results describe a process of late postnatal development of top-down mPFC afferents onto DRN causally linked to the emergence of behavioral persistence in the transition between adolescence and adulthood. This critical period of corticofugal axonal development may also represent a period of vulnerability for maladaptive development involved in the etiopathogenesis of psychiatric disorders (Rutter, 2007; Chen et al., 2019; Soiza-Reilly et al., 2019; Guirado et al., 2020).

## Methods

### Animals

All experimental procedures were approved and performed in accordance with the Champalimaud Centre for the Unknown Ethics Committee guidelines and by the Portuguese Veterinary General Board (Direcção-Geral de Veterinária, approval 0421/000/000/2016). The mouse lines used in this study were obtained from the Mutant Mouse Resource and Research Center (MMRRC), Rbp4-Cre (stock number 031125-UCD), and from Jax Mice, Ai32(RCL-ChR2(H134R)/EYFP) (Stock number 012569) and Ai9(RCL-tdTomato) (7905). All of them were crossbred in-house for at least 10 generations prior to their use in our experiments. Mice were kept under a standard 12 h light/dark cycle with food and water ad libitum. Behavioral testing occurred during the light period.

### Electrophysiological recordings

Male and female mice were used for whole-cell recordings. Coronal slices of 300 µm thickness containing the dorsal raphe were cut using a vibratome (Leica VT1200) in “ice cold” solution containing (in mM): 2.5 KCl, 1.25 NaH2PO4, 26 NaHCO3, 10 D-glucose, 230 Sucrose, 0.5 CaCl2, 10 MgSO4, and bubbled with 5% CO2 and 95% O2. Slices were recovered in ACSF containing (in mM): 127 NaCl, 2.5 KCl, 25 NaHCO3, 1.25 NaH2PO4, 25 Glucose, 2 CaCl2, 1 MgCl2 at 34 °C for 30 minutes and then kept in the same solution at room temperature until transferred to the recording chamber. In addition, 300uM L-Tryptophan (Sigma) was added to the ACSF to maintain serotonergic tone in the ex vivo preparation as described elsewhere (Liu et al., 2005).

Patch recording pipettes (resistance 3-5 MΩ) were filled with internal solution containing (in mM): 135 K-Gluconate, 10 HEPES, 10 Na-Phosphocreatine, 3 Na-L-Ascorbate, 4 MgCl2, 4 Na2-ATP, and 0.4 Na-GTP. Data were acquired using a Multiclamp 700B amplifier and digitized at 10 kHz with a Digidata 1440a digitizer (both from Molecular Devices). Data was then either analyzed using Clampfit 10.7 Software (Molecular Devices, LLC) or imported into Matlab and analyzed with custom-written software.

Every voltage-clamp recording contained a 100ms test pulse of -10mV for offline calculation of access and series resistance to ensure the same recording quality across experiments. The access resistance (Ra) was determined by measuring the amplitude of the current response to the command voltage step and the membrane resistance (Rm) as the difference between the baseline and the holding current in the steady state after the capacitive decay, by applying Ohm’s law. Input resistance was the sum of the membrane resistance with the pipette resistance. The membrane time constant (τ) was determined by a single exponential fit of the decay phase in response to the square pulse. An approximation of the capacitance was obtained using the following formula:

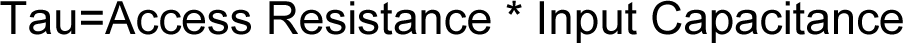

Only neurons with access resistance 1/10th lower than the membrane resistance were used for analysis. Only neurons with an access resistance lower than 30MOhm were considered for analysis. The access resistance was comparable between neurons recorded at different developmental stages (ANOVA, F(3,119)=1.78, p=0.15). Neurons were recorded at a holding membrane voltage of -70mV, near the reversal potential of chloride (-68mV) and thus, optogenetically evoked responses correspond to AMPA-mediated currents.

To activate ChR2-expressing fibers, light from a 473-nm fiber-coupled laser (PSU-H-FDA, CNI Laser) was delivered at approximately 2mm from the sample to produce wide-field illumination of the recorded cell (Fig. 2A). TTL triggered pulses of light (10-ms duration; 10 mW measured at the fiber tip, which was located approximately 2 mm away from the sample) were delivered at the recording site at 0.1Hz of frequency. Neurons were recorded in the dorsomedial and lateral wings portions of the DRN, in two consecutive coronal slices per mouse between bregma levels ∼4.3 and ∼4.8 (Fig. S1).

To assess ChR2 evoked photocurrent in layer V somas, mPFC slices were obtained using the same slicing procedure. 1 µM TTX was added to the bath to prevent escaped spikes in the voltage clamp recordings upon light activation.

### Histology

Mice were deeply anesthetized with pentobarbital (Eutasil) and perfused transcardially with 4% paraformaldehyde (P6148, Sigma-Aldrich). The brain was removed from the skull, stored in 4% paraformaldehyde for two hours before being transferred to cryoprotectant solution (30% sucrose in PBS) until they sank. Sagittal sections (50 µm) were cut with a freezing sliding microtome (SM2000, Leica).

For axonal quantification in Rbp4-ChR2, we performed anti-GFP immunostaining to enhance the intrinsic signal of the ChR2-fused EYFP reporter. We incubated overnight with the anti-GFP primary antibody at 4 degrees (1:1000, A-6455 Invitrogen, 0.1M PBS 0.3% tx100, 3% NGS). After abundant PBS washes, we incubated the secondary biotinylated anti-Rabbit antibody (711-065-152, Jackson IRL) for 2-4 hours in the same incubation solution at room temperature and finally, after PBS washes, slices were incubated in Alexa488 Streptavidin for 2-4 hours in the same incubation solution at room temperature (S32354, Invitrogen). After final PBS washes, slices were mounted and covered with FluoroGel mounting medium (17985-10, Electron Microscopy Sciences) for posterior imaging.

### Image acquisition and analysis

Histological sections were imaged with a Zeiss LSM 710 confocal laser scanning microscope using 10x and 25x magnification objectives.

To quantify the axonal density, images containing the DRN or the nucleus accumbens were background subtracted and binarized using constant thresholds in Fiji. After thresholding, binary images were imported into Matlab and analyzed with custom written software. Images were sampled every 100um across the Y axis. The intersections between binary axons and these sampling lines across the Y axis were counted and averaged in bins of 100um to estimate axonal density.

Quantification of layer V soma densities was obtained in histological slices of Rbp4-tdTomato mice. Confocal images were imported into matlab, and fluorescent somas were detected using the image analysis toolbox of Matlab inside a defined region of interest containing the mPFC (PL/IL). The number of somas was then divided by the area of the ROI to obtain the density of neurons.

### Stereotaxic surgeries and virus injection

Animals were anesthetized with isoflurane (2% induction and 0.5 - 1% for maintenance) and placed in a motorized computer-controlled Stoelting stereotaxic instrument with mouse brain atlas integration and real-time surgery probe visualization in the atlas space (Neurostar, Sindelfingen, Germany;https://www.neurostar.de). Antibiotic (Enrofloxacin, 2.5-5 mg/Kg, S.C.), pain killer (Buprenorphine,0.1 mg/Kg, S.C.), and local anesthesia over the scalp (0.2 ml, 2%Lidocaine, S.C.) were administered before incising the scalp. Virus injection (experiment group: AAV2retro-flex-EF1A-taCasp3-TEVp; control group: AAV2retro-flex-hSyn-tdTomato) was targeted to DRN at the following coordinates: -4.7 mm AP, 0.0 mm ML, and 3.1 mm DV. The vertical stereotaxic arm was tilted 32 degrees caudally to reach the target avoiding Superior sagittal sinus and Transverse sinuses. Target coordinates were adjusted as follows: -6.64 mm AP, 0.0 mm ML, and -4.02 mm DV. To infect a larger volume of the DRN with the virus, we performed six injections of 0.2 uL using two entry points along the AP axis (-6.54 and -6.74) and 3 depths along the DV axis(-4.02, -3.92, and -3.82). The incision was then closed using tissue adhesive (VETBOND^TM/MC^, 3M, No. 1469SB). Mice were monitored until recovery from the surgery and returned to their homecages, where they were housed individually. Behavioral testing started at least 1 week after surgery to allow for recovery.

### Behavioral testing

The behavioral box consisted of 1 back-wall (16x219 cm), 2 side-walls (16.7x219 cm), and 2 front-walls (10x219 cm,140-degrees angle between them), made of white acrylic (0.5 cm thick) and a transparent acrylic lead. A camera (ELP camera, ELP-USBFHD01M-L180) was mounted on top of the ceiling for monitoring purposes. Each front wall had a nose-poke port equipped with an infrared emitter/sensor pairs to report port entry and exit times (model 007120.0002, Island motion corporation) and a water valve for water delivery (LHDA1233115H, The Lee Company, Westbrook, CT). An internal white acrylic wall (8cm) separates the two nose-poke ports forcing the animals to walk around it to travel between ports. All signals from sensors were processed by Arduino Mega 2560 microcontroller board (Arduino, Somerville, US), and outputs from the Arduino Mega 2560 microcontroller board were implemented to control water delivery in drops of 4μl. Arduino Mega 2560 microcontroller was connected to the sensors and controllers through an Arduino Mega 2560 adaptor board developed by the Champalimaud Foundation Scientific Hardware Platform.

Subjects have to probe two foraging sites (nose-poke ports, for mice, or virtual magic wands, for humans) to obtain rewards (4 μl water drops, for mice, or virtual points for humans). At any given time, only one of the sites is active and, when probed, delivers a reward with a fixed 90% probability (PREW). Each attempt also triggers a fixed 90% probability of transition (PTRS) to inactivate the current foraging site and activate the other. These transitions are not cued; thus, subjects are required to alternate probing the current site and traveling to the other to track the hidden active state and obtain rewards. In this work, we focus on assessing differences in the baseline patience/impulsivity, measured as the ability to withhold adverse outcomes. Therefore naive subjects were only tested once.

Five days before testing, water dispensers were removed from the animals’ home cages, and their weights were recorded. In the following days, progressively less water (1000μl, 800μl, 600μl) was given in a metal dish inside the homecage. Weight loss was monitored every day before water delivery, and no animal lost more than 20% of their body weight. On the fifth day of water deprivation, animals were weighed and introduced to the behavioral box. A small quantity of water was present at the start of the session to stimulate the mice to probe the nose-ports. Sessions lasted a minimum of one hour. By that time, if animals did not perform at least 30 trials, the session was extended for thirty more minutes.

Animals were handled during water deprivation to reduce stress levels, but they were completely naive about the task environment and functioning on the testing day. One juvenile female mouse was excluded from the experiment batch before the task assessment because of congenital blindness. One caspase adult mouse was excluded after the task assessment because of abnormal behavior. Rather than nose poking to seek water, this animal spent most of the task time biting the nose port, to anomalous levels. In chronological order, we tested a batch of only male juvenile and adult animals, followed by testing of male and female tdTomato and Caspase animals, and finally only female juvenile and adult animals. Separate analysis for females and males on the effect of age reveals that juveniles are less persistent in both cases.

### Data and statistical analysis

Behavioral data analysis was performed using custom-written scripts in Julia-1.4.1. Behavioral results were represented as median ± 95% confidence intervals, and statistical significance was accepted for p-values< 0.05. The statistical analysis was done in Julia-1.4.1 (Bezanson et al., 2017) with the HypothesisTests (https://juliastats.org/HypothesisTests.jl/v0.9/) and MixedModels (Bates et al., 2021) existing packages. The effect of a specific factor on the probability of leaving was tested by applying logistic regression on a generalized linear mixed-effects model (GLMM), using a Bernoulli distribution for the dependent variable and a Logit link function. For each foraging nose poke we assigned a boolean label according to whether the animal left the patch after that poke (True) or not (False). We then use logistic regression to explain this leaving choice for each poke according to the elapsed time in the trial (PokeTime), the elapsed trials in the session (Trial), the animal group (Age or Virus) and their interactions. This statistical approach allows us to examine the question of behavioral persistence in terms of probability of leaving after each single poke, expanding the amount of usable data, per animal and counterbalancing the limitation of studying the phenomenon in naive animals exposed to a single session. Furthermore this technique can test for both additive and multiplicative effects of the factors contributing to behavioral persistence. The individual variability was accounted for through generalized linear mixed models with random intercept and slopes for each mouse (see methods for the implementation). Before testing we checked for co-linearity between the continuous predictors and confirmed that there was no correlation between the time of poking (Poke Time) and trials elapsed from the beginning of the session (Trial) (PokeTime∼1+Trial+(1+Trial|MouseID): p = 0.99, Fig, S3B). First, to assess the significance of the estimated coefficients, we calculated their 95% CI by performing a parametric bootstrap of 1000 samples. Only factors whose CI did not include 0 were considered to be significantly affecting the probability of leaving. Next, to validate the relevance of the experimental manipulation (age or virus), we compared nested models (a general model and a special case model, excluding or including the experimental factor, respectively) using a likelihood ratio test: chi-squared test on the difference of the deviance of the two nested models, with degrees of freedom equal to the difference in degrees of freedom between the general model (lacking the predictor) and its special case (with the predictor of interest). For each analysis, we report the median and 95% CI of the median for the groups of interest, followed by the test statistics. We use Wilkinson annotation to describe the models with denoting random effects.

Electrophysiological and histological results were analyzed with Matlab and Graphpad Software. Normality of the residuals was tested with the D’Agostino-Pearson omnibus K2 test. When normally distributed, either a t-test, one-way ANOVA or repeated measures ANOVA were performed to compare groups at different developmental phases. In the cases where residuals were not normally distributed, we performed a Mann-Whitney or Kruskal Wallis test to assess significance. For testing differences in connection probability, a Chi-square test was performed. Finally, a Kolmogorov Smirnoff test was performed to compare the neuronal density distribution between Caspase treated animals and tdTomato expressing controls. Error bar plots represent mean ± SEM. Significance was noted as *p<0.05.

## Acknowledgements

We thank Drs. Cindy Poo and Constanze Lenschow for helpful comments on the manuscript and the Champalimaud Foundation Advanced Bio-optics and Bio-imaging platform for the microscopy technical assistance. This work was supported by the Champalimaud Foundation (Z.F.M.), European Research Council (671251, Z.F.M.), Fundação para a Ciência e Tecnologia (FCT-PTDC/MED-NEU/28830/2017, Z.F.M.; SFRH / BD / 132172 / 2017, D.S.). This work was further supported by Portuguese national funds Fundação para a Ciência e a Tecnologia (FCT; UIDB/04443/2020); CONGENTO, co-financed by Lisboa Regional Operational Programme (Lisboa2020), under the PORTUGAL 2020 Partnership Agreement, through the European Regional Development Fund (ERDF) and Fundação para a Ciência e Tecnologia (Portugal) under the project LISBOA-01-0145-FEDER-022170, the imaging platform has been financed under the project LISBOA-01-0145-FEDER-022122.

## Author contributions

NGC, DS and ZFM designed the research. NGC, DS and BSG performed the research. NGC and DS analyzed the data. NGC, DS and ZFM wrote the paper.

## Supplementary Figures

**Supplementary Figure 1.**
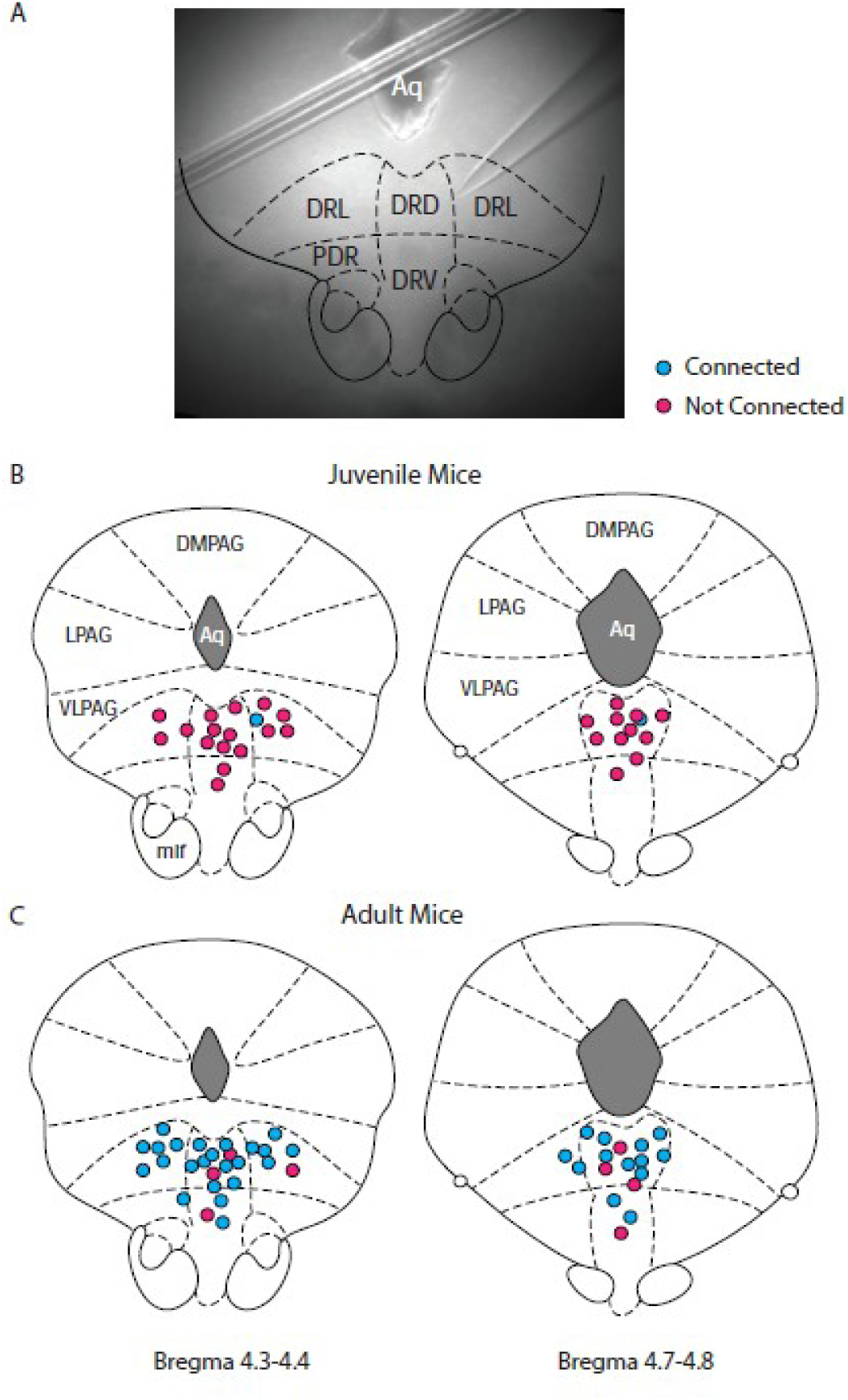
Changes in cortico-raphe connectivity over development are not explained by changes in the location of the recorded DRN neurons. (A) Example low magnification picture taken of a recorded DRN neuron and overlaid atlas inset used to determine its location. (B,C) Summary of the spatial location and connectivity of the recorded DRN neurons in juvenile (anterior DRN connected/non-connected = 1/16, posterior DRN connected/non-connected = 1/11) and adult (anterior DRN connected/non-connected=23/4, posterior DRN connected/non-connected = 13/4) mice.

**Supplementary Figure 2.**
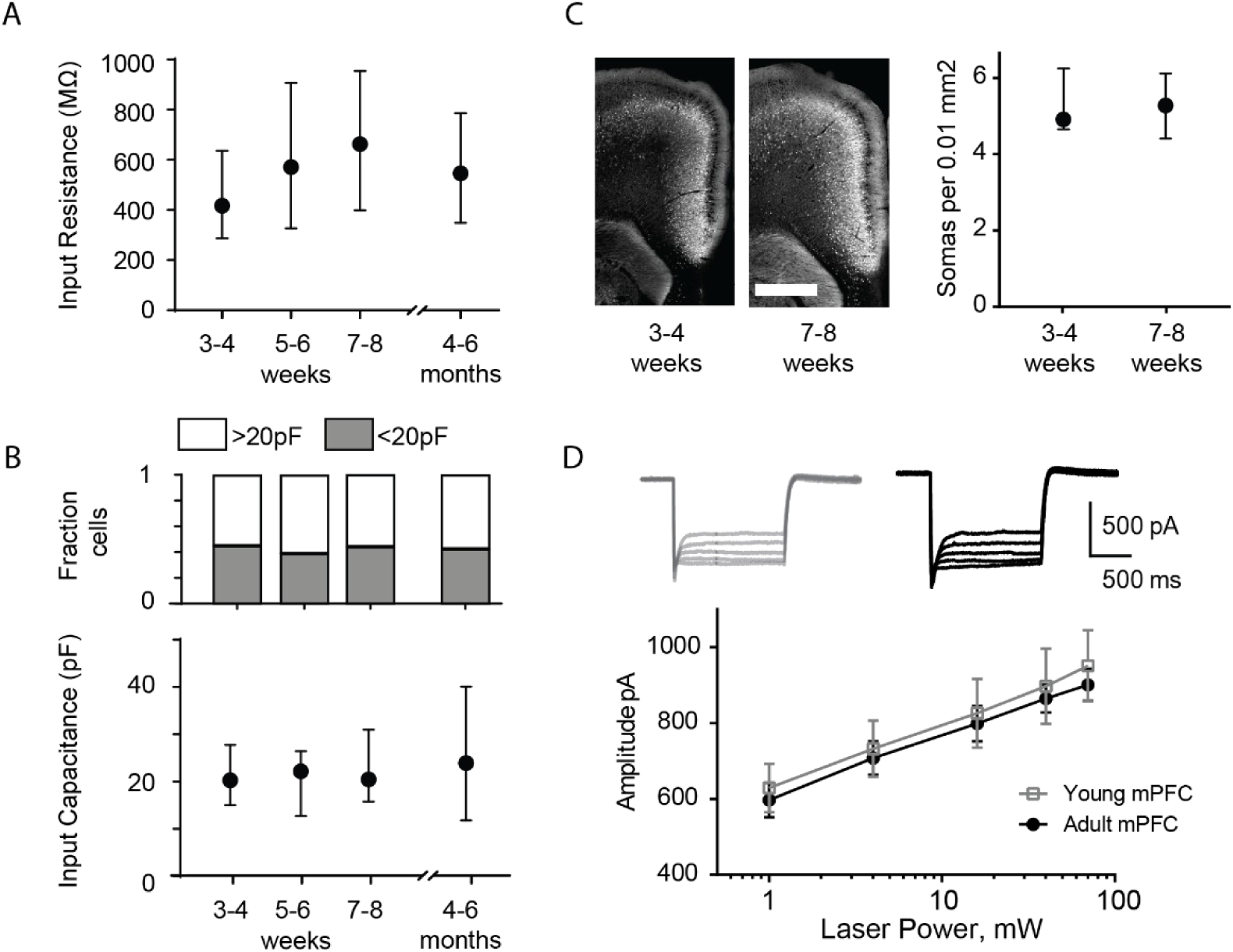
Changes in cortico-raphe connectivity over development are not explained by changes in membrane properties of DRN neurons or by differential ChR2 expression of ChR2 under the Rbp4 promoter over time. (A) The input resistance of DRN neurons is comparable over time. (B) The fraction of putative 5HT neurons (Capacitance>20pF) and non-5HT neurons (Capacitance<20pF) (Soiza-Reilly et al., 2019) recorded is comparable across developmental stages (Fraction of neurons with Capacitance>20pF: 3-4 weeks= 0.55, 5-6 weeks= 0.62, 7-8 weeks= 0.54, 5-6 months= 0.6. Chi-Square test Χ^2^ (3, N=122 neurons) =0.59, p=0.89)). In addition, no overall changes in input capacitance were observed in DRN neurons across development. (C) The density of fluorescent mPFC layer V neurons is comparable in juvenile and adult Rbp4-tdTomato mice. (D) The evoked photocurrent in mPFC layer V neurons of juvenile and adult Rbp4-ChR2 mice is virtually identical across a wide range of stimulation intensities. Error bars in A-C represent median and 95% CI. Error bars in D represent mean ± SEM. Scale bar in C = 800 microns.

**Supplementary Figure 3.**
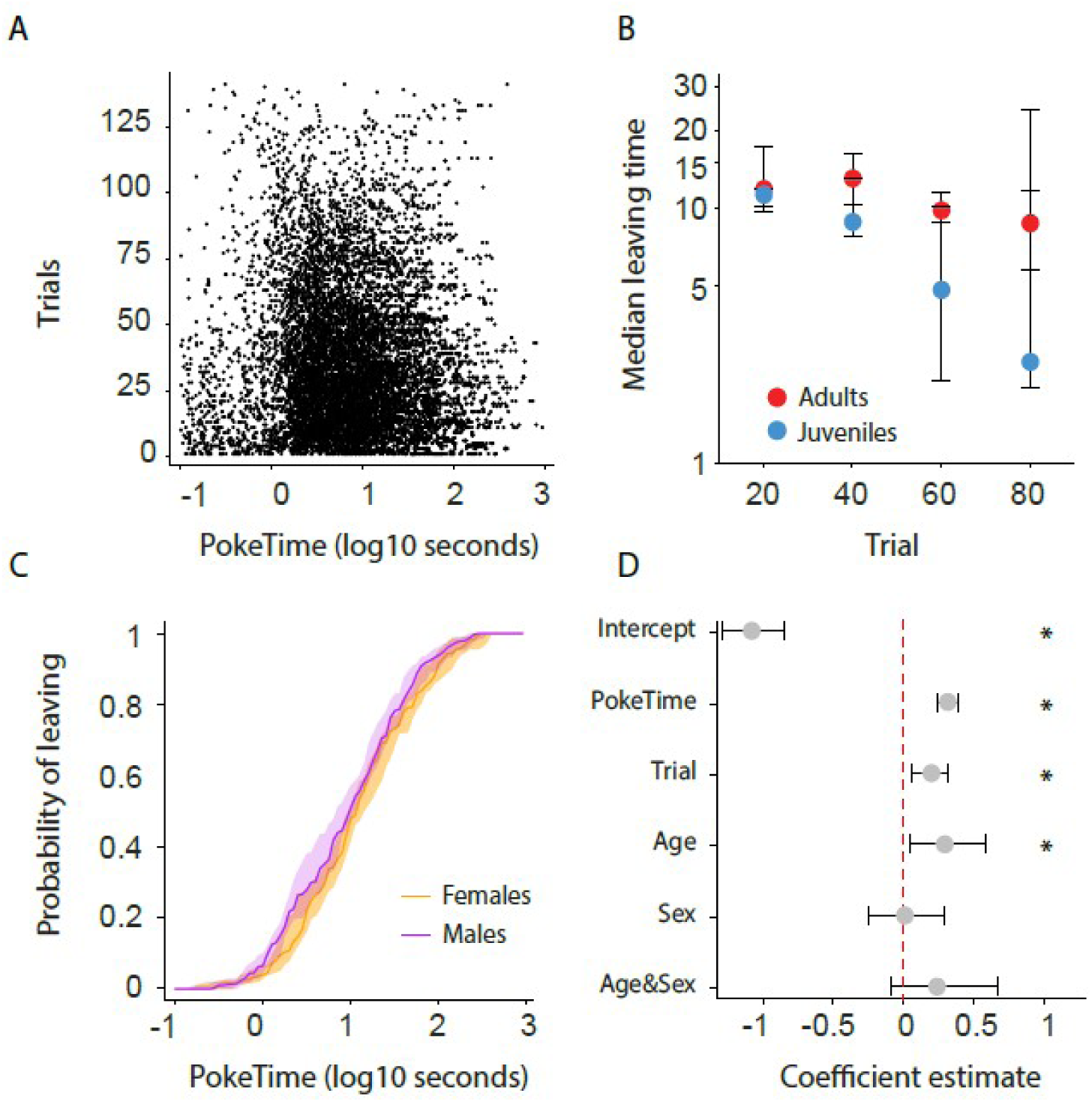
Description of poking behavior over the session progression and according to sex. (A) Individual poke durations for all mice. (B) Leaving time (median ± 95% CI across mice) as a function of elapsed trials in a session. (C) Cumulative distribution of the probability of leaving as a function of trial time elapsed (median ± 95% CI across mice) for female and male mice. (D) Regression coefficients ± 95% CI resulting from a parametric bootstrap (n = 1000) of a mixed models logistic regression to explain the probability of leaving. Note the lack of explanatory power for the group variable sex.

**Supplementary Figure 4.**
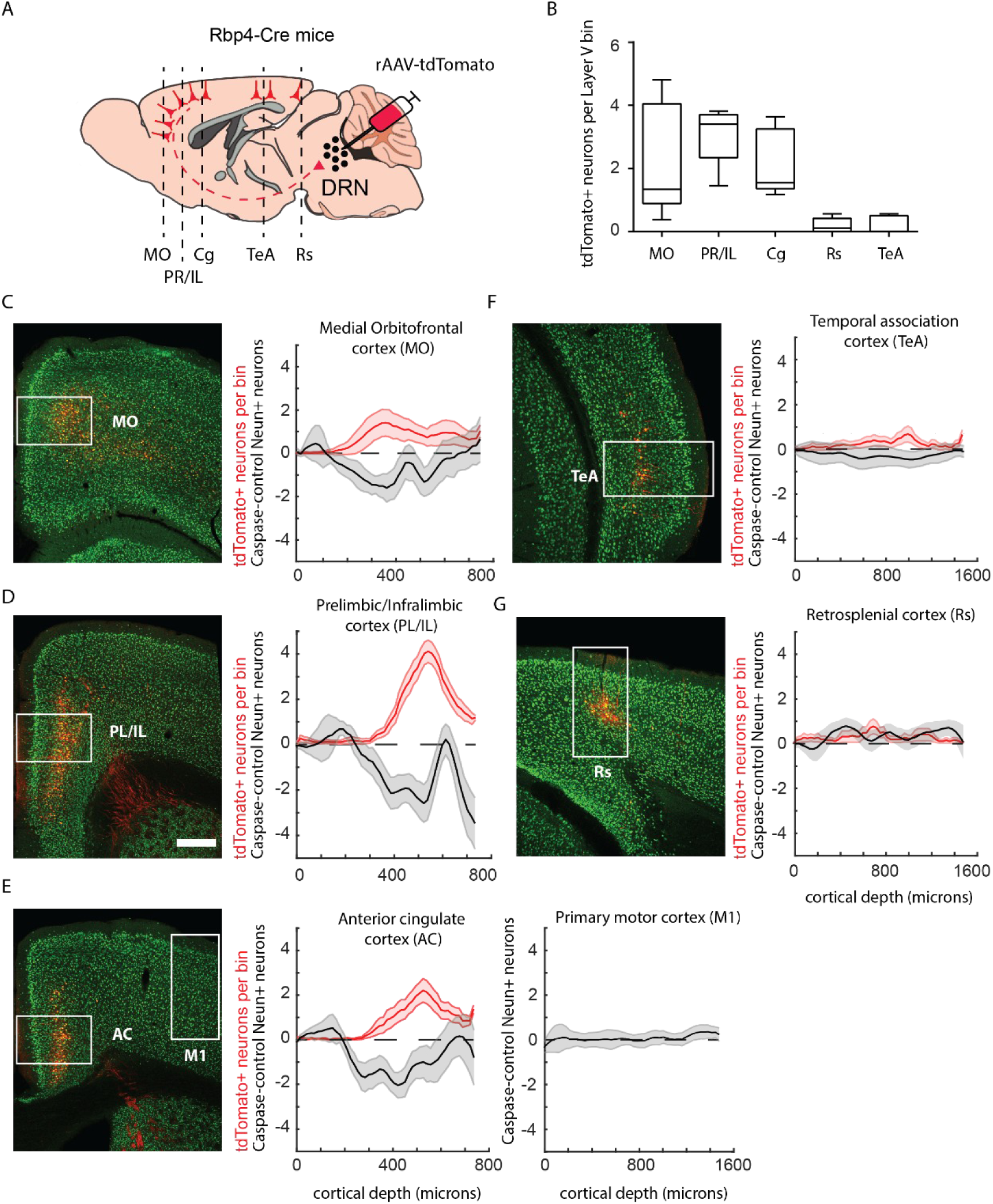
Labeling of Rbp4-expressing DRN projecting neurons with rAAV-tdTomato is consistent with cell density loss in mice injected with rAAV-Caspase3. (A) Schematic representation of rAAV-tdTomato dependent labeling of cortico-DRN projecting neurons in Rbp4-Cre mice. (B) Quantification of layer V tdTomato labeled somas across the different DRN projecting cortical areas (n = 8 mice). Example picture of the immunolabeling obtained with the pan-neuronal marker NeuN and the virally expressed tdTomato reporter together with the quantification across cortical depth of tdTomato cell density and neuronal loss (NeuN density in rAAV-Caspase3 injected mice - average NeuN density of control mice, n=7 caspase mice) for the medial orbitofrontal cortex (C, Control vs. Caspase Two-sample Kolmogorov-Smirnoff Test, D = 0.017, p = 0.08), prelimbic/infralimbic cortex (D, D = 0.028, p = 0.002), cingulate cortex and motor primary cortex (E, D = 0.024, p = 0.01 and D = 0.019, p = 0.15, respectively), temporal association cortex (F, D = 0.025, p = 0.19) and retrosplenial cortex (G, D = 0.034, p = 0.12). Box plots represent median, IQR, and min/max data range. Shaded error plots represent mean ± SEM. Scale bar = 400 microns.

**Supplementary Figure 5.**
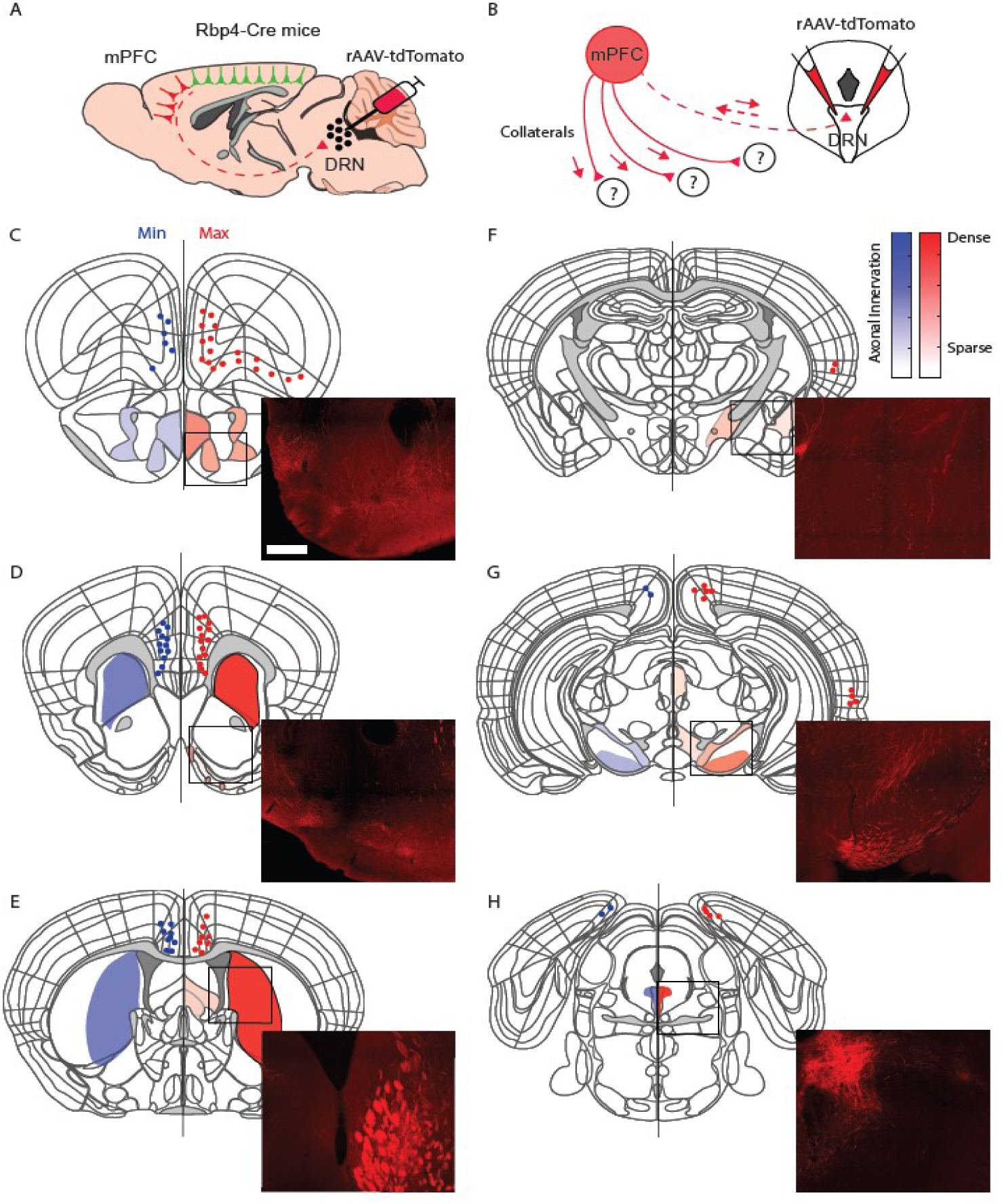
Dorsal raphe projecting cortical neurons have dense collateral projections to the striatum. (A) Schematic representation of rAAV-tdTomato dependent labeling of cortico-DRN projecting neurons in Rbp4-Cre mice. (B) Schematic representation of axon collaterals from the same cortical subpopulation of neurons retrogradely labeled at the DRN. (C-H) Semiquantitative representation of axon collateral innervation density across the anteroposterior levels of the mouse brain presenting the injections with the highest (red) and lowest (blue) density of retrogradely labeled neurons. Scale bar= 500 microns.

